# Comparative Pathogenesis Of COVID-19, MERS And SARS In A Non-Human Primate Model

**DOI:** 10.1101/2020.03.17.995639

**Authors:** Barry Rockx, Thijs Kuiken, Sander Herfst, Theo Bestebroer, Mart M. Lamers, Dennis de Meulder, Geert van Amerongen, Judith van den Brand, Nisreen M.A. Okba, Debby Schipper, Peter van Run, Lonneke Leijten, Ernst Verschoor, Babs Verstrepen, Jan Langermans, Christian Drosten, Martje Fentener van Vlissingen, Ron Fouchier, Rik de Swart, Marion Koopmans, Bart L. Haagmans

**Affiliations:** Department of Viroscience, Erasmus University Medical Center, Rotterdam, the Netherlands; Viroclinics Xplore, Schaijk, the Netherlands; Department of Virology, Biomedical Primate Research Centre, Rijswijk, the Netherlands; Institute of Virology, Charité-Universitätsmedizin, Berlin, Germany; Erasmus Laboratory Animal Science Center, Erasmus University Medical Center, Rotterdam, the Netherlands

## Abstract

A novel coronavirus, SARS-CoV-2, was recently identified in patients with an acute respiratory syndrome, COVID-19. To compare its pathogenesis with that of previously emerging coronaviruses, we inoculated cynomolgus macaques with SARS-CoV-2 or MERS-CoV and compared with historical SARS-CoV infections. In SARS-CoV-2-infected macaques, virus was excreted from nose and throat in absence of clinical signs, and detected in type I and II pneumocytes in foci of diffuse alveolar damage and mucous glands of the nasal cavity. In SARS-CoV-infection, lung lesions were typically more severe, while they were milder in MERS-CoV infection, where virus was detected mainly in type II pneumocytes. These data show that SARS-CoV-2 can cause a COVID-19-like disease, and suggest that the severity of SARS-CoV-2 infection is intermediate between that of SARS-CoV and MERS-CoV.

**One Sentence Summary:** SARS-CoV-2 infection in macaques results in COVID-19-like disease with prolonged virus excretion from nose and throat in absence of clinical signs.

---

Following the first reports of an outbreak of an acute respiratory syndrome (2019 coronavirus disease, or COVID-19) in China in December 2019, a novel coronavirus, SARS-CoV-2, was identified (*1, 2*). As of March 14, 2020, over 140,000 cases were reported worldwide with over 5,400 deaths, surpassing the combined number of cases and deaths of two previously emerging coronaviruses, SARS-CoV and MERS-CoV (*3*). COVID-19 is characterized by a range of symptoms, including fever, cough, dyspnoea and myalgia in most cases (*2*). In severe cases, bilateral lung involvement with ground-glass opacity is the most common chest CT finding (*4*). Similar to the 2002/2003 outbreak of SARS, severity of COVID-19 disease is associated with increased age and/or a comorbidity, although severe disease is not limited to these risk groups (*5*). However, despite the large number of cases and deaths, limited information is available on the pathogenesis of this virus infection. Two reports on the histological examination of the lungs of three patients showed bilateral diffuse alveolar damage (DAD), pulmonary oedema and hyaline membrane formation, indicative of acute respiratory distress syndrome (ARDS), as well as characteristic syncytial cells in the alveolar lumen (*6, 7*), similar to findings during the 2002/2003 outbreak of SARS-CoV (*8*). The pathogenesis of SARS-CoV infection was previously studied in a non-human primate model where aged animals were more likely to develop disease (*9–13*). In the current study, SARS-CoV-2 infection was characterized in the same animal model, using cynomolgus macaques, and compared with infection with MERS-CoV and historical data on SARS-CoV (*9, 10, 12*).

First, two groups of four cynomolgus macaques (both young adult, 4-5 years of age; and aged, 15-20 years of age) were inoculated via a combined intratracheal (IT) and intranasal (IN) route with a SARS-CoV-2 strain from a German traveller returning from China. No overt clinical signs were observed in any of the infected animals, with the exception of a serous nasal discharge in one aged animal on day 14 post inoculation (p.i.). No significant weight loss was observed in any of the animals during the study. By day 14 p.i. all remaining animals seroconverted as revealed by the presence of SARS-CoV-2 specific antibodies against the S1 domain and nucleocapsid proteins in their sera (Fig. S1).

As a measure of virus shedding, nasal, throat and rectal swabs were assayed for virus by RT-qPCR and virus culture. In nasal swabs, detection of SARS-CoV-2 RNA peaked by day 2 p.i. in young adult animals and day 4 p.i. in aged animals and was detected up to at least day 8 p.i. in two out of four animals and up to day 21 p.i. in one out of four animals (Fig. 1A). Overall, higher levels of SARS-CoV-2 RNA were detected in nasal swabs of aged animals compared to young adult animals. SARS-CoV-2 RNA detection in throat swabs peaked at day 1 p.i. in young adult and day 4 p.i. in aged animals and decreased more rapidly over time, compared to nasal swabs, but could still be detected intermittently up to day 10 p.i. (Fig. 1B). Low levels of infectious virus could be cultured from nasal and throat swabs up to day 4 and 2 respectively (Table S1). SARS-CoV-2 RNA could only be detected in a rectal swab from 1 animal on day 14 p.i. and no viral RNA could be detected in whole blood at any time point throughout the study.

**Fig. 1.**
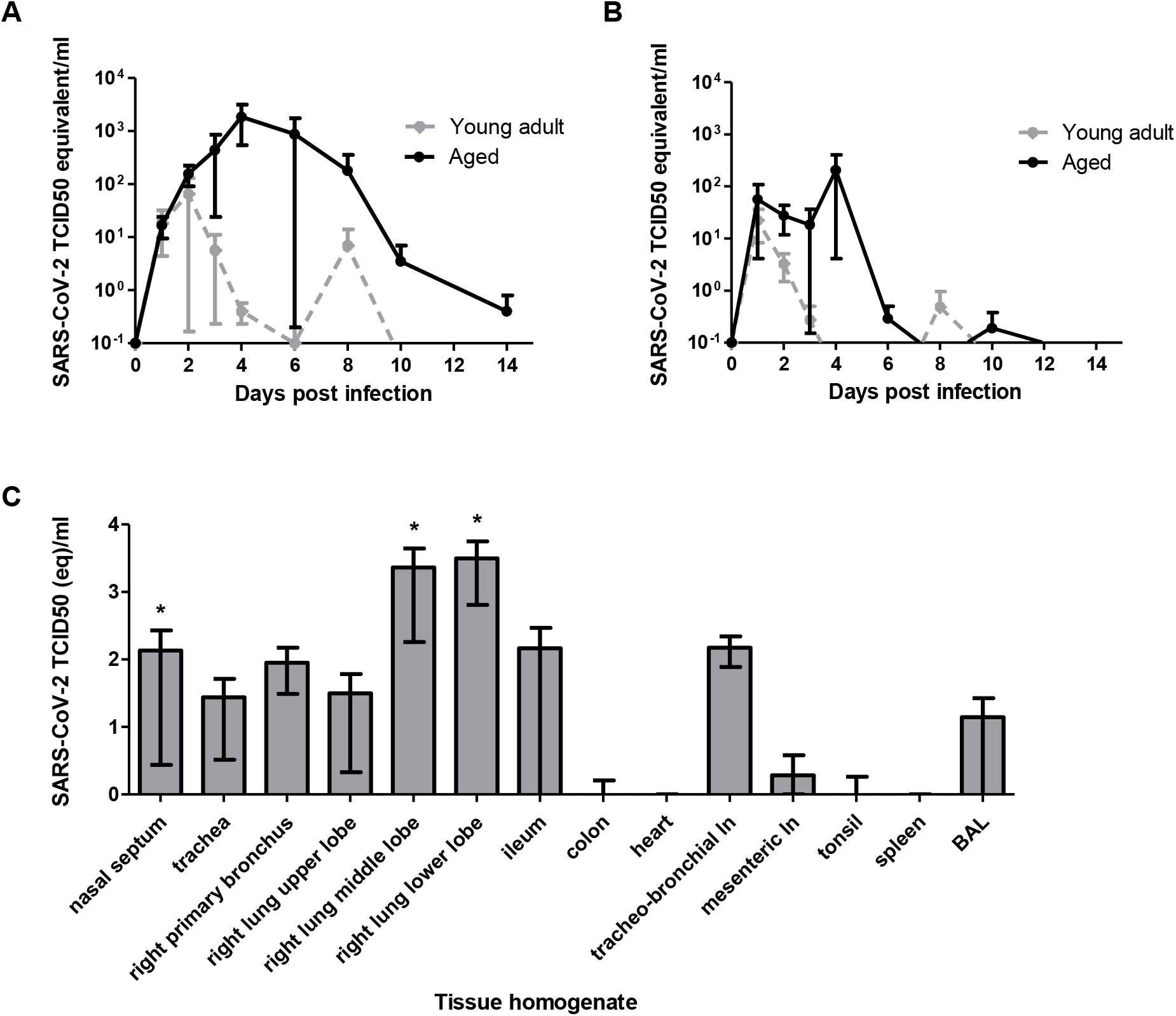
Virus shedding and virus detection in organs of SARS-CoV-2 inoculated cynomolgus macaques. Viral RNA was detected in nasal (A) and throat (B) swabs and tissues (C) of SARS-CoV-2 infected animals by RT-qPCR. Samples from 4 animals (days 1-4) or 2 animals (days >4) per group were tested. The error bars represent the standard error of the mean. Virus was detected in tissues from 2 young, and 2 aged animals on day 4 by RT-qPCR.* = infectious virus was isolated.

Upon autopsy of four macaques on day 4 p.i., two had foci of pulmonary consolidation in the lungs (Fig. 2A). One animal (aged: 17 years) in the right middle lobe, representing less than 5% of the lung tissue. A second animal (young adult: 5 years) had two foci in the left lower lobe, representing about 10% of the lung tissue (Fig. 2A). The consolidated lung tissue was well-circumscribed, red-purple, level, and less buoyant than normal. The other organs in these two macaques, as well as the respiratory tract and other organs of the other two animals were normal.

**Fig. 2.**
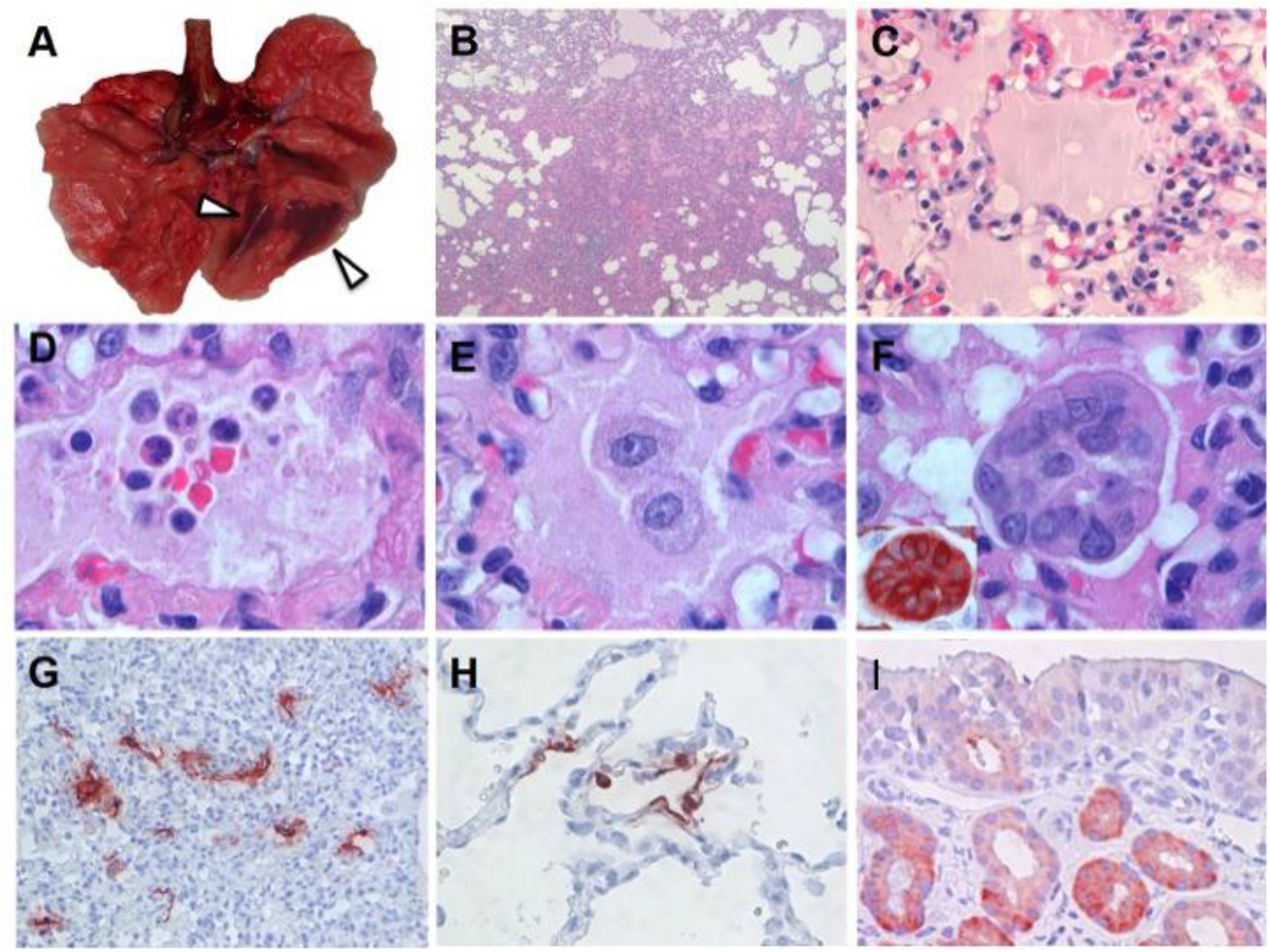
Characteristic pathological changes and virus antigen expression in the lungs of SARS-CoV-2-inoculated cynomolgus macaques. (**A**) Two foci of pulmonary consolidation in the left lower lung lobe (arrowheads). (**B**) Area of pneumonia (HE, 2X objective). (**C**) Oedema fluid in alveolar lumina (HE, 40X objective). (**D**) Neutrophils, as well as erythrocytes, fibrin, and cell debris, in an alveolar lumen flooded by oedema fluid (HE, 100X objective). (**E**) Mononuclear cells, either type II pneumocytes or alveolar macrophages, in an alveolar lumen flooded by oedema fluid (HE, 100X objective). (**F**) Syncytium in an alveolar lumen (HE, 100X objective). Inset: Syncytium expresses keratin, indicating epithelial cell origin (IHC for pankeratin AE1/AE3, 100X objective). (**G**) SARS-CoV-2 antigen expression is colocalized with areas of diffuse alveolar damage (IHC for SARS-CoV-nucleocapsid, 20X objective). (**H**) Type I (flat) and type II (cuboidal) pneumocytes in affected lung tissue express SARS-CoV-2 antigen (IHC for SARS-CoV-nucleocapsid, 40X objective). (**I**) Epithelial cells of mucous glands in nasal cavity expresses SARS-CoV-2 antigen (IHC for SARS-CoV-nucleocapsid, 40X objective).

Virus replication was assessed by RT-qPCR on day 4 p.i. in tissues from the respiratory, digestive urinary, and cardiovascular tracts, from endocrine and central nervous systems, as well as from various lymphoid tissues. Virus replication was primarily restricted to the respiratory tract (nasal cavity, trachea, bronchi and lung lobes) with highest levels of SARS-CoV-2 RNA in lungs (Fig. 1C). Interestingly, in three out of four animals, SARS-CoV-RNA was also detected in ileum and tracheo-bronchial lymph nodes (Fig. 1C).

The main histological lesion in the consolidated pulmonary tissue involved the alveoli and bronchioles, and consisted of areas with acute or more advanced DAD (Fig. 2B). In these areas, the lumina of alveoli and bronchioles were variably filled with protein-rich oedema fluid, fibrin, and cellular debris, a moderate number of alveolar macrophages, and fewer neutrophils and lymphocytes (Fig. 2C-E). There was epithelial necrosis with extensive loss of epithelium from alveolar and bronchiolar walls. Hyaline membranes were present in a few damaged alveoli. In areas with more advanced lesions, the alveolar walls were moderately thickened and lined by cuboidal epithelial cells (type II pneumocyte hyperplasia), and the alveolar lumina were empty. Alveolar and bronchiolar walls were thickened by oedema fluid, mononuclear cells, and neutrophils. There were aggregates of lymphocytes around small pulmonary vessels. Moderate numbers of lymphocytes and macrophages were present in the lamina propria and submucosa of the bronchial walls, and a few neutrophils in the bronchial epithelium. Regeneration of epithelium was seen in some bronchioles, visible as an irregular layer of squamous to high cuboidal epithelial cells with hyperchromatic nuclei. There were occasional multinucleated giant cells (syncytia) free in the lumina of bronchioles and alveoli (Fig. 2F), and, based on positive pan-keratin staining and negative CD68 staining, originated from epithelial cells (Fig. 2F, inset).

SARS-CoV-2 antigen expression was detected in moderate numbers of type I pneumocytes and a few type II pneumocytes in foci of DAD (Fig. 2G, fig. S2). The pattern of staining was similar to that in lung tissue from SARS-CoV-infected macaques (positive control). SARS-CoV-2 antigen expression was not observed in any of the syncytia. In the upper respiratory tract, there was focal or locally extensive SARS-CoV-2 antigen expression in epithelial cells of mucous glands in the nasal cavity (septum or concha) of all four macaques, without any associated histological lesions (fig. 2I).

In order to assess the severity of infection with SARS-CoV-2 compared to that with the previously emerged MERS-CoV, young adult cynomolgus macaques (3-5 years of age) were inoculated with MERS-CoV via the IN and IT route. All animals remained free of clinical signs. At day 21 p.i., all remaining animals (n=2) seroconverted as revealed by the presence of MERS-CoV specific antibodies in their sera by ELISA.

MERS-CoV RNA was detected in nasal (Fig. 3A) and throat swabs (Fig. 3B) from days 1 to 11 p.i., with peaks on days 1 and 2 p.i., respectively. Low levels of MERS-CoV RNA were detected in rectal swabs on days 2 and 3 p.i. (data not shown).

**Fig. 3.**
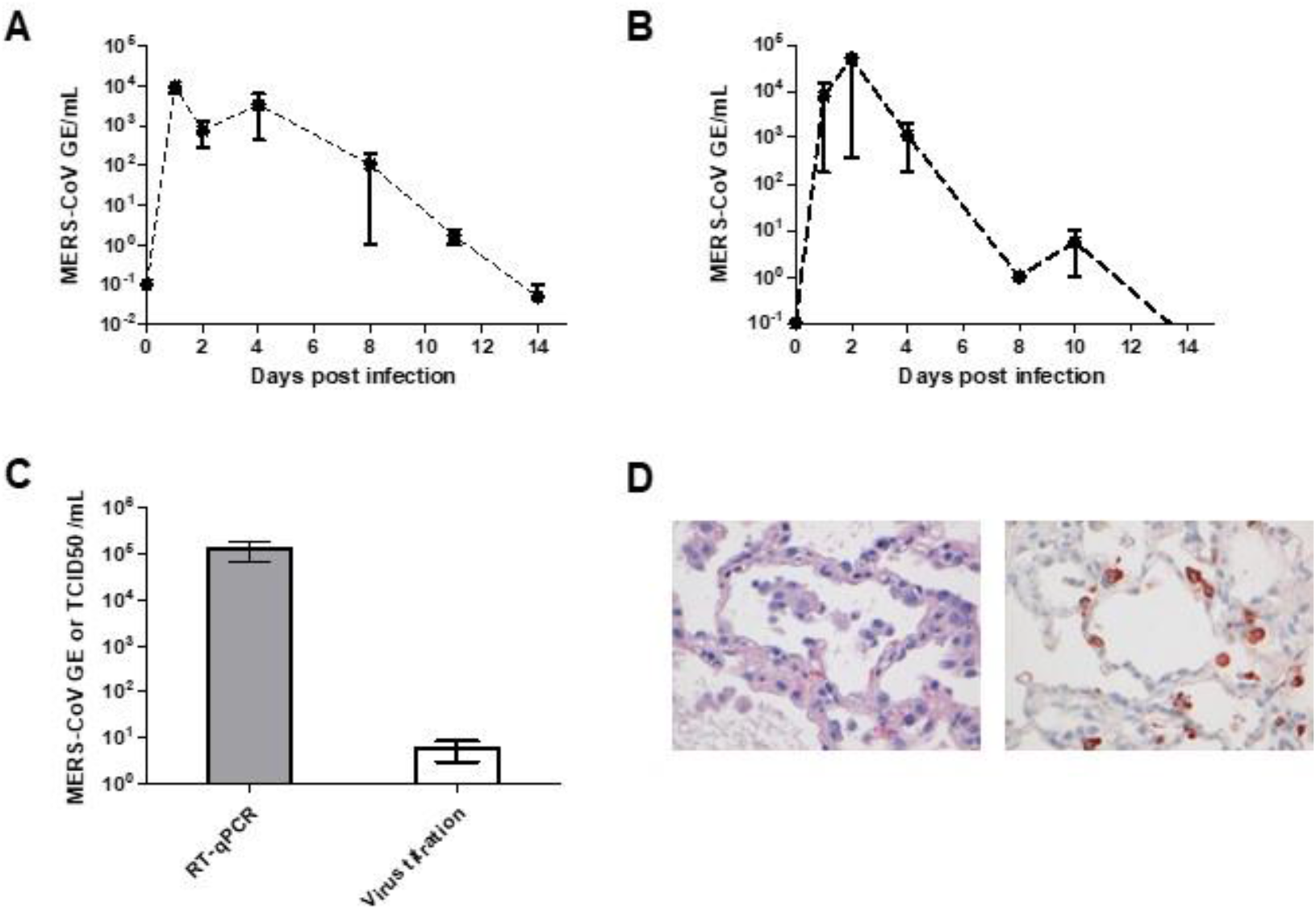
Virus shedding and virus detection in organs of MERS-CoV inoculated cynomolgus macaques. Viral RNA was detected in nasal (A) and throat (B) swabs and tissues (C) of MERS-CoV infected animals by RT-qPCR. Samples from 4 animals per group were tested. The error bars represent the standard error of the mean. Virus was detected in tissues on day 4 by RT-qPCR. Histopathological changes (D; left picture) with hypertrophic and hyperplastic type II pneumocytes in the alveolar septum and increased numbers of alveolar macrophages in the alveolar lumen and virus antigen expression (right picture) in type II pneumocytes.

Upon autopsy of four macaques at day 4 p.i., three out of four macaques had foci of pulmonary consolidation, characterized by slightly depressed areas in the lungs, representing less than 5 % of the lung tissue (Table 1). Similar to SARS-CoV-2, on day 4 p.i., MERS-CoV RNA was primarily detected in the respiratory tract of inoculated animals (Fig. 3C). Infectious virus titres were comparable to SARS-CoV-2, but lower compared to SARS-CoV infection of young adult macaques (Table 1). In addition, MERS-CoV RNA was detected in the spleen (Table 1).

**Table 1.** Comparative pathogenesis of SARS-CoV-2, MERS-CoV, and SARS-CoV infections in cynomolgus macaques.

| Virus | Age category | No. | Clinical signs | Maximum excretion from throat | Maximum excretion from nose | Viral titre in lung | Virus in extra-respiratory tissues | Pulmonary lesions at 4 days post inoculation |  |  |  |  | Reference |
| --- | --- | --- | --- | --- | --- | --- | --- | --- | --- | --- | --- | --- | --- |
|  |  |  |  |  |  |  |  | % affected lung | Hyaline membranes | Alveolar oedema | Leukocyte infiltration | Cell type tropism |  |
| **SARS-CoV** | Young adult | 4 | No | 10e3.5 TCID50eq | 10e4.2 TCID50eq | 10e6.5 TCID50eq | No | 0 - 5 | No | No | Yes | Type I & II pneumocytes | (10-12) |
| **SARS-CoV** | Aged | 4 | Yes | 10e3.0 TCID50eq | 10e3.0 TCID50eq | 10e6.2 TCID50eq | Kidney | 0 - 60 | Yes | Yes | Yes | Type I & II pneumocytes | (10-12) |
| **MERS-CoV** | Young adult | 4 | No | 10e5.3 TCID50eq | 10e4.3 TCID50eq | 10e4.4 TCID50eq | Spleen | 0 - 5 | No | Yes (small amount) | Yes | Type II pneumocytes | This manuscript |
| **SARS-CoV-2** | Young adult | 2 | No | 10e1.8 TCID50eq | 10e2.4 TCID50eq | 10e1.1 TCID50eq | Ileum, colon, tonsil | 0 – 10 | Yes | Yes | Yes | Type I & II pneumocytes | This manuscript |
| **SARS-CoV-2** | Aged | 2 | No | 10e2.9 TCID50eq | 10e3.7 TCID50eq | 10e0.7 TCID50eq | Ileum, colon, tonsil | 0 - 5 | No | No | Yes | Type I & II pneumocytes | This manuscript |

Consistent with the presence of virus in the lower respiratory tract at day 4 p.i., histopathological changes characteristic for DAD were observed in the lungs of inoculated animals (Fig. 3D). The alveolar septa were thickened due to infiltration of few neutrophils and macrophages, and moderate type II pneumocyte hyperplasia and hypertrophy. In the alveolar lumina, there were increased numbers of alveolar macrophages and occasionally small amounts of oedema fluid with fibrin and few neutrophils (Fig. 3D). Few syncytial cells were seen in the alveolar lumina. Interestingly, MERS-CoV antigen was not detected in tissues on day 4 p.i. in any part of the respiratory tract. Therefore, four young adult macaques were included at an earlier time point, day 1 p.i. On day 1 p.i. there was multifocal expression of viral antigen predominantly in type II pneumocytes and occasionally in type I pneumocytes, bronchiolar and bronchial epithelial cells and few macrophages (Fig 3D).

In summary, we inoculated young adult and aged cynomolgus macaques with a low passage clinical isolates of SARS-CoV-2, which resulted in productive infection in the absence of overt clinical signs. Increased age did not affect disease outcome, but there was prolonged viral shedding in the upper respiratory tract of aged animals. Prolonged shedding has been observed in both SARS-CoV-2 and SARS-CoV patients (*14, 15*). Interestingly, SARS-CoV-2 in our model, peaked early in the course of infection, similar to what is seen in patients (*14*). This earlier peak in virus shedding for SARS-CoV-2 is more similar to influenza virus shedding (*16*) than SARS-CoV, where shedding peaked around day 10 post onset of symptoms (*15*), and may explain why case detection and isolation may not be as effective for SARS-CoV-2 as it was for the control of SARS-CoV (*17*). Also, all four macaques expressed SARS-CoV-2 antigen in mucous glands of the nasal cavity at day 4 p.i., which was not seen for SARS-CoV *(9-11)* or MERS-CoV infections (this manuscript) in this animal model. Viral tropism for the nasal mucosa fits with efficient respiratory transmission, as has been seen for influenza A virus (*18*). SARS-CoV-2 was primarily detected in tissues of the respiratory tract, however SARS-CoV-2 RNA was also detectable in other tissues such as intestines, in line with a recent report (*19*).

Two out of four animals had foci of DAD on day 4 p.i. The colocalisation of SARS-CoV-2-antigen expression and DAD provides strong evidence that SARS-CoV-2 infection caused this lesion. The histological character of the DAD, including alveolar and bronchiolar epithelial necrosis, alveolar oedema, hyaline membrane formation, and accumulation of neutrophils, macrophages and lymphocytes, corresponds with the limited pathological analyses of human COVID-19 cases (*6, 7*). In particular, the presence of syncytia in the lung lesions is characteristic of respiratory coronavirus infections. Whereas MERS-CoV primarily infects type II pneumocytes in cynomolgus macaques, both SARS-CoV and SARS-CoV-2 also infect type I pneumocytes. Injury to type I pneumocytes can result in pulmonary oedema, and formation of hyaline membranes (*20*), which may explain why hyaline membrane formation is a hallmark for SARS and COVID-19 (*7, 10*), but not frequently reported for MERS (*21, 22*).

These data show that cynomolgus macaques are permissive to SARS-CoV-2 infection, shed virus for a prolonged period of time and display COVID-19-like disease. SARS-CoV-2 infection caused histopathological changes in the lungs that appeared to be intermediate in severity between those from SARS-CoV and MERS-CoV infections established by similar inoculation doses and routes. An in-depth comparison of infection with SARS-CoV, MERS-CoV and SARS-CoV-2 in this model may identify key pathways in the pathogenesis of these emerging viruses. This study provides a novel infection model which will be critical in the evaluation and licensure of preventive and therapeutic strategies against SARS-CoV-2 infection for use in humans, as well as evaluating the efficacy of repurposing species specific existing treatments, such as pegylated interferon (*12*).

## Acknowledgments

We thank L. de Meulder, A. van der Linden, I. Chestakova and R. Sikkema for technical assistance. Y.Kap, D. Akkermans, V. Vaes, M. Sommers, F. Meijers for assistance with the animal studies.

## Author contributions

Conceptualization, B.R. R.F., M.K., RdS, M.F. B.H.; investigation, B.R., T.K., S.H., T.B., M.L., D.d.M., G.v.A., J.v.d.B., N.O., D.S., P.v.R., L.L.; resources, B.H., C.D. E.V., B.V., J.L. supervision, B.R. and B.H.; writing, original draft, B.R., T.K. and B.H.; writing–review and editing, all authors; funding acquisition: B.H., R.F. and M.K.

## Data and materials availability

All data is available in the main text or the supplementary materials.

