## Supplemental Materials for "Comparative Pathogenesis Of COVID-19, MERS And SARS In A Non-Human Primate Model"

**This PDF file includes:**

Materials and Methods
Figs. S1 to S2
Tables S1

### Materials and Methods

Viruses and cells

SARS-CoV-2 (isolate BetaCoV/Munich/BavPat1/2020) was obtained from a clinical case in Germany diagnosed after returning from China (European Virus Archive Global # 026V-03883). The virus was propagated to passage 3 on Vero E6 cells in Opti-MEM I (1X) + GlutaMAX (Gibco), supplemented with penicillin (10,000 IU/mL) and streptomycin (10,000 IU/mL) at 37°C in a humidified CO2 incubator. MERS-CoV (EMC strain, accession no. NC_019843; European Virus Archive Global # 011V-02838), was grown as previously described (*1*). All work was performed in a Class II Biosafety Cabinet under BSL-3 conditions at Erasmus Medical Center.

Animals and Ethical Statement

Healthy, purpose bred, adult female cynomolgus macaques (*Macaca fascicularis*) were handled in an ABSL3 biocontainment laboratory. Research was conducted in compliance with the Dutch legislation for the protection of animals used for scientific purposes (2014, implementing EU Directive 2010/63) and other relevant regulations. The licensed establishment where this research was conducted (Erasmus MC) has an approved OLAW Assurance # A5051-01. Research was conducted under a project license from the Dutch competent authority and the study protocol was approved by the institutional Animal Welfare Body. All steps were taken to minimize the number of animals and to protect their welfare, in particular to avoid any unnecessary suffering of the animals in accordance with the ‘‘Weatherall report for the use of non-human primates’’ recommendations, and early clinical humane endpoint criteria applied. Animals were housed in pairs in primate cages in Class III isolators allowing social interactions, under controlled conditions of humidity, temperature and light (12-hour light/12-hour dark cycles). Food and water were available ad libitum and commercial monkey chow was supplemented with fresh fruits for enrichment. Animals were cared for and monitored (pre- and post-infection) by qualified personnel. The animals were sedated/anesthetized for all invasive procedures.

Animal procedures SARS-CoV-2

Eight female cynomolgus macaques (ages 5–20 years) weighing between 3,5 and 5,0 kg were distributed evenly regarding age over two groups of four animals. Animals were seropositive for herpesvirus B. Each group consisted of two young adult animals (5 years) and two aged animals (15-20 years). Animals were inoculated with SARS-CoV-2 under anesthesia via a combination of intratracheal (4.5 ml) and intranasal (0.25 ml per nostril) routes with a suspension containing 2x10e5 TCID50 per ml PBS (total infectious dose = 10e6 TCID50). Animals were anesthetized for challenge, blood collection, and swabs of nasal, throat and rectal mucosa on days 0, 1, 2, 3, 4, 6, 8, 10, 14, 18 and 21 p.i. One group of four animals (two young adult and 2 aged) was euthanized and autopsied on day 4 p.i. with collection of tissue specimens from respiratory, digestive urinary, and cardiovascular tracts, endocrine and central nervous systems, as well as various lymphoid organs. The other group was euthanized at day 21 p.i. with collection of tissue specimens from respiratory, digestive urinary, and cardiovascular tracts, endocrine and central nervous systems, as well as various lymphoid organs.

### Animal procedures MERS-CoV

Ten female young adult cynomolgus macaques (3-5 years) weighing between 3-4 kg were distributed evenly regarding age over two groups of four animals and one group of two animals. Animals were inoculated with MERS-CoV under anesthesia through a combination of intratracheal (4.5 ml) and intranasal (0.25 ml per nostril) inoculation with a suspension containing 2x10e5 TCID50 per ml PBS (total infectious dose = 10e6 TCID50). Animals were anesthetized for challenge, blood collections, and taking nasal, throat and rectal swabs on days 0, 1, 2, 3, 4, 6, 8, 10, 14, 18 and 21 p.i. Groups of four animals were euthanized and autopsied on days 1 and 4 p.i. with collection of tissue specimens from respiratory, digestive urinary, and cardiovascular tracts, endocrine and central nervous systems, as well as various lymphoid organs. One group of 2 animals was euthanized on day 21 p.i.

### Serological Analysis

To test for SARS-CoV-2 antibodies, serum samples were collected at days 0, 4, 8, 14, and 21. Serum samples were tested for SARS-CoV-2 antibodies using an spike S1 and nucleocapsid (N) ELISAs (manuscript in preparation). Briefly, ELISA plates were coated overnight with either SARS-CoV-2 S1 or SARS-CoV N proteins. After blocking, serum samples were added and incubated for 1h at 37$^{o}$C. Bound antibodies were detected using HRP-labeled rabbit anti–human IgG (Dako) and TMB (Life Technologies) as a substrate. The absorbance of each sample was measured at 450 nm.

### Virus detection

Samples from nasal septum, trachea, right primary bronchus, right upper, middle and lower lung lobe, bronchoalveolar lavage (right upper lung lobe), heart, liver, pancreas, stomach, duodenum, jejunum, ileum, colon, kidney, urinary bladder, uterus, ovary, cerebrum, cerebellum, brain stem, olfactory bulb, tracheo-bronchial and mesenteric lymphnodes, tonsil and spleen were collected post mortem for virus detection by RT-qPCR and virus isolation as previously described for MERS-CoV and SARS-CoV (*2-4*). Briefly, tissues were homogenized 10% w/v in viral transport medium using Polytron PT2100 tissue grinders (Kinematica). After low-speed centrifugation, the homogenates were frozen at −70°C until they were inoculated on Vero E6 cell cultures in 10-fold serial dilutions. The SARS-CoV-2 RT-qPCR was performed as previously published (*5*). The MERS-CoV RT-qPCR was performed as previously published (*2*).

Pathology

Autopsies of the animals were performed according to a standard protocol. For histological examination the following tissues were collected: adrenal gland, aorta, axillary lymph node, brachial biceps muscle, brain stem, caecum, cerebellum, cerebrum, colon, duodenum, eye, eyelid, femoral bone marrow, heart muscle (left and right), ileum, inguinal lymph node, jejunum, kidney, lung (inflated with 10% neutral-buffered formalin), liver, mandibular lymph node, mesenteric lymph node, nasal concha, nasal septum, oesophagus, palatum molle, pancreas, primary bronchus, ovary, salivary gland, spleen, stomach, thymus, thyroid gland, tongue, tonsil, trachea, tracheobronchial lymph node, urinary bladder, and uterus. Tissues for light-microscope examination were fixed in 10% neutral-buffered formalin, embedded in paraffin, and 3 µm sections were stained with haematoxylin and eosin.

Immunohistochemistry

SARS-CoV-2 and MERS-CoV antigens were detected by immunohistochemistry in duplicate sections of all tissue samples. Paraffin was removed from sections, which were rehydrated and pretreated with citric acid buffer (pH 6.0) for 15 min at 100°C. Endogenous peroxidase was blocked with 3% hydrogen peroxide. Slides were briefly washed with 0,05% phosphate-buffered saline Tween 20 (Fluka, Buchs, Switzerland) and blocked with 10% goat serum (X0907, DAKO, Agilent Technologies Netherlands B.V, Amstelveen, the Netherlands) for 30 min at room temperature. Slides were then incubated with a rabbit polyclonal antibody against SARS-CoV-nucleoprotein (40143-T62, Sino Biological, Chesterbrook, PA, USA) or Rabbit IgG isotypecontrol (AB-105-C, R&D, Minneapolis, MN, USA) diluted 1:1000 in phosphate-buffered saline with 0,1% bovine serum albumin for 1 h at room temperature. After washing, sections were incubated with horseradish peroxidase labeled goat-anti-rabbit IgG (P0448, DAKO, Agilent Technologies Netherlands B.V. Amstelveen, The Netherlands), diluted 1:100 in phosphate-buffered saline with 0,1% bovine serum albumin for 1h at room temperature. Horseradish peroxidase activity was revealed by incubating slides in 3-amino-9-ethylcarbazole (Sigma, St Louis, MO, USA) solution for 10 min, resulting in a bright red precipitate. Sections were counterstained with haematoxylin. MERS-CoV antigen was detected with a rabbit-anti-SARS-CoV NSP4 which cross-reacts with MERS-CoV (*2*). Historical tissue sections from cynomolgus macaques that had not been infected with SARS-CoV-2 were included as negative controls. Historical tissue sections from cynomolgus macaques that had been infected with SARS-CoV were included as positive controls.

For staining of epithelial cells, slides (processed as above) were incubated with mouse-anti-human pankeratin AE1/AE3 IgG1 (Neomarkers, Fremont, CA, USA) (2 µg/ml) in phosphate-buffered saline with 0,1% bovine serum albumin for 1h at room temperature. To detect macrophages, slides were incubated with mouse-anti-human CD68 IgG1 (clone KP1, DAKO) (1:200) in phosphate-buffered saline with 0,1% bovine serum albumin for 1h at room temperature. As negative control, mouse IgG1 isotype control (R&D Systems Europe, Abingdon, UK ) (2 µg/ml) was included. After washing, all sections were incubated with horseradish peroxidase labeled goat-anti-mouse IgG1 (Southern Biotech, Birmingham, AL, USA) in phosphate-buffered saline with 0,1% bovine serum albumin for 1 h at room temperature. Horseradish peroxidase activity was revealed by incubating slides in 3-amino-9-ethylcarbazole (Sigma) solution for 10 min, resulting in a bright red precipitate, followed by counterstaining with haematoxylin.

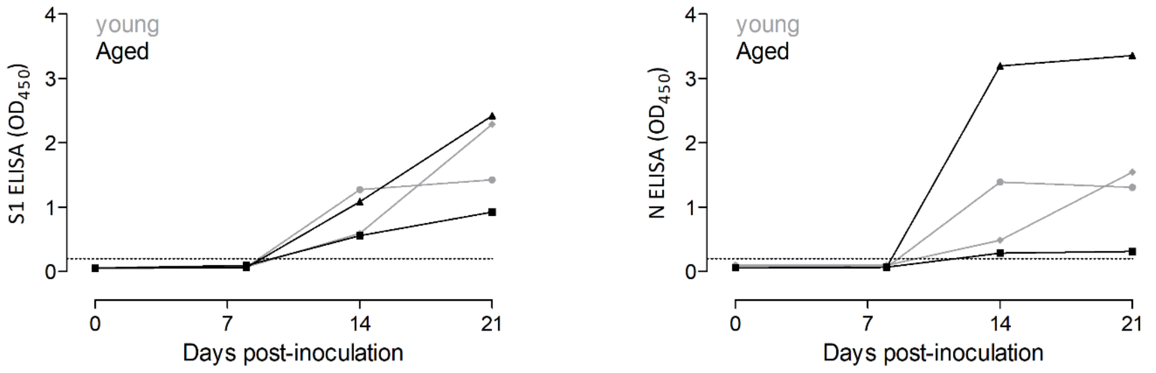


**Fig. S1.**
SARS-CoV-2 specific antibody responses in inoculated cynomolgus macaques. Serum was collected from four SARS-CoV-2 inoculated cynomolgus macaques, and IgG responses were evaluated using an SARS-CoV-2 recombinant spike protein domain 1 (S1) and nucleoprotein ELISA. Sera were assayed on days 0, 8, 14 and 21 p.i., and the optical density at 450nm is shown on the y axis. Sera from young adult animals are shown in gray, sera from aged animals are shown in black.

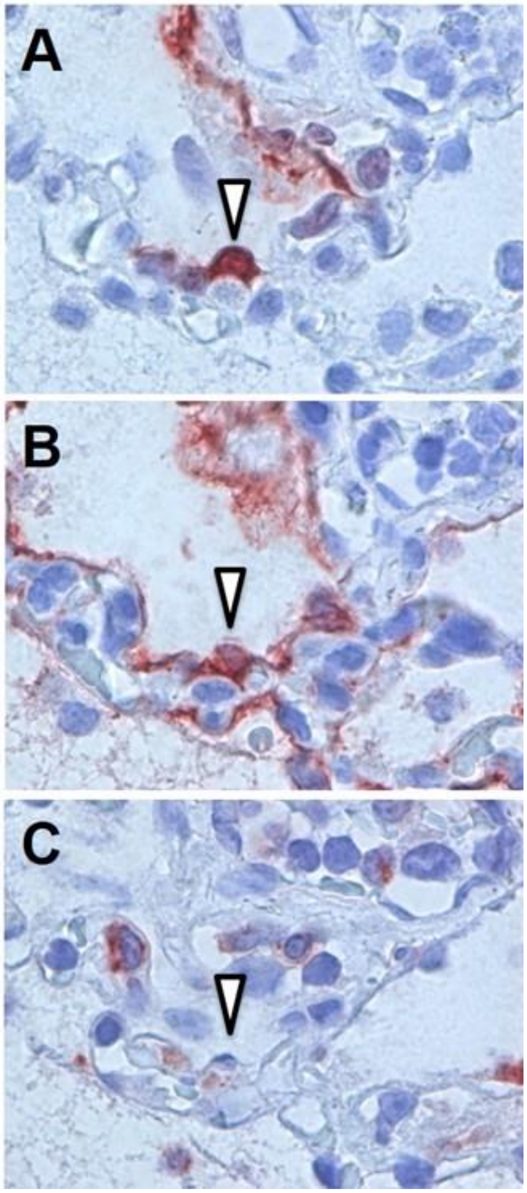


**Fig. S2.**

Characteristics of cuboidal cells expressing SARS-CoV-2 antigen in lung of macaque inoculated with SARS-CoV-2. (**A**) Cuboidal cell lining alveolar wall expresses SARS-CoV-2 antigen (IHC for SARS-CoV-nucleocapsid). (**B**) In sequential tissue section, cell at same location as in A expresses keratin, indicating it is a type II pneumocyte (IHC for pankeratin AE1/AE3). (**C**) In sequential tissue section, cell at same location as in A does not express CD68, indicating it is not an alveolar macrophage (IHC for CD68). All panels: 100X objective.

**Table S1.**
Detection of SARS-CoV-2 RNA and infectious virus in nasal and throat swabs.

| | Nasal swabs | | Throat swabs | |
|---|---|---|---|---|
| **Days post inoculation** | **RT-qPCR** | **Virus culture** | **RT-qPCR** | **Virus culture** |
| 0 | 0/8 | 0/8 | 0/8 | 0/8 |
| 1 | 7/8 | 0/8 | 7/8 | 4/8 |
| 2 | 7/8 | 2/8 | 8/8 | 1/8 |
| 3 | 7/8 | 1/8 | 8/8 | 0/8 |
| 4 | 7/8 | 1/8 | 4/8 | 0/8 |
| 6 | 3/4 | 0/4 | 2/4 | 0/4 |
| 8 | 2/4 | 0/4 | 1/4 | 0/4 |
| 10 | 1/4 | ND | 1/4 | ND |
| 14 | 1/4 | ND | 0/4 | ND |
| 18 | 1/4 | ND | 1/4 | ND |
| 21 | 1/4 | ND | 1/4 | ND |
